# Morphogenesis of neurons and glia within an epithelium

**DOI:** 10.1101/393850

**Authors:** Isabel I. C. Low, Claire R. Williams, Megan K. Chong, Ian G. McLachlan, Bradley M. Wierbowski, Irina Kolotuev, Maxwell G. Heiman

## Abstract

To sense the outside world, some neurons protrude across epithelia, the cellular barriers that line every surface of our bodies. To study the morphogenesis of such neurons, we examined the *C. elegans* amphid, in which dendrites protrude through a glial channel at the nose. During development, amphid dendrites extend by attaching to the nose via DYF-7, a type of protein typically found in epithelial apical ECM. Here, we show that amphid neurons and glia exhibit epithelial properties, including tight junctions and apical-basal polarity, and develop in a manner resembling other epithelia. We find that DYF-7 is a fibril-forming apical ECM component that prevents rupture of the tube-shaped glial channel, reminiscent of roles for apical ECM in other narrow epithelial tubes. We also identify a role for FRM-2, a homolog of EPBL15/moe/Yurt which promote epithelial integrity in other systems. Finally, we show that other environmentally-exposed neurons share a requirement for DYF-7. Together, our results suggest that these neurons and glia can be viewed as part of an epithelium continuous with the skin, and are shaped by mechanisms shared with other epithelia.

## INTRODUCTION

Between each of us and the outer world sits an epithelium. These sheets of cells, held together by tight and adherens junctions, create a barrier at every surface of our body and provide the interface through which we perceive our surroundings.

Neurons form intimate connections with epithelia in order to transmit information from the exterior surfaces of the body to the brain. For example, our sense of touch is mediated by mechanosensory neurons that branch along the basal (“inner”) side of the skin epithelium. The skin plays a major role in shaping these sensory neurons, and these epithelia-neuron interactions have been extensively studied in vertebrates (Wang et al., 2013; Zimmerman et al., 2014), *Drosophila* (Parrish et al., 2009; Han et al., 2012; Kim et al., 2012; Jiang et al., 2014; Tenenbaum et al., 2017), and *C. elegans* (Dong et al., 2013; Liang et al., 2015; Dĺaz-Balzac et al., 2016; Liu et al., 2016; Zou et al., 2016; Zhu et al., 2017; Celestrin et al., 2018). Similarly, our senses of hearing and taste are mediated by afferent neurons that are positioned at the basal surfaces of epithelia and receive information from specialized non-neuronal sensory cells (hair cells, taste cells) within these epithelia (Roper, 2013; Frank and Goodrich, 2018). In each of these examples, neurons make intimate connections with epithelia, but they remain restricted to the “inside” surface and do not directly contact the outside world.

In contrast, our sense of smell is mediated by sensory neurons with a different topology: they protrude through the epithelium to the apical (“outside”) surface where they detect odorants in the outside world (Firestein, 2001). On one side, the neurons extend dendrites that form tight junctions with glial-like cells in the epithelium (Steinke et al., 2008). On the other side, the neurons extend axons that pierce the basal lamina of the epithelium and extend into the brain (Graziadei and Graziadei, 1979). Thus, based on their anatomy, these sensory neurons have properties of both epithelial cells and neurons – like an epithelial cell, they have an outward facing surface exposed to the environment but, like a true neuron, they have an axon that projects into the brain. Sensory neurons with this anatomy are prevalent in invertebrates, and might represent an ancestral sensory structure (McLachlan and Heiman, 2013). In most cases, they are surrounded by glial supporting cells that are also exposed to the outside environment. These interesting structures raise several questions. Do the neurons and glia exhibit other epithelial-like properties, such as apical-basal polarity? Do they develop using mechanisms of morphogenesis that are similar to those used by epithelia? What role do the glia play during neuron development?

Here, we address these questions using the *C. elegans* amphid, which consists of 12 neurons and two glial cells (Ward et al., 1975). Each neuron extends an unbranched dendrite to the nose where it senses environmental stimuli, as well as an axon that projects into the nerve ring, or “brain” (White et al., 1986). The two glial cells, called the sheath and socket, each extend a single process to the nose where they form a tube-shaped pore through which most of the dendrites have direct access to the outside environment (Ward et al., 1975). The sheath completely wraps the dendrite endings, creating a local environment that can affect sensory function (Perens and Shaham, 2005; Bacaj et al., 2008; Wang et al., 2008; Oikonomou et al., 2011; Procko et al., 2011; Oikonomou et al., 2012; Singhvi et al., 2016; Wallace et al., 2016; Wang et al., 2017), while the socket forms an opening continuous with the skin. The dendrite endings protrude through this tube-shaped glial pore into the environment, and are decorated with cilia bearing receptors for specific smells, tastes, or other stimuli (Bargmann, 2006).
In the embryo, the amphid dendrites develop by a mechanism called retrograde extension, which is distinct from the typical growth-cone-mediated outgrowth of axons or dendrites (Heiman and Shaham, 2009). First, amphid neurons and glia are assembled into a polarized multicellular rosette that is carried to the nose by the migrating skin epithelium (Fan et al., 2018). Then, the nascent tips of the amphid dendrites stay anchored at the nose while the neuronal cell bodies migrate away, stretching the dendrites out behind them (Heiman and Shaham, 2009). DYF-7, a zona pellucida (ZP) domain protein, is required to keep the dendrite tips anchored at the nose; in the absence of DYF-7, the nascent dendrites are dragged along behind the migrating neuronal cell bodies and fail to extend (Heiman and Shaham, 2009). Interestingly, ZP domain proteins like DYF-7 are typically associated with epithelia, rather than with developing neurons (Plaza et al., 2010). They are almost always found in the apical extracellular matrix (ECM), which coats the outer (or luminal) surfaces of epithelia in the kidney, liver, and many other tissues. ZP domain proteins, along with other apical ECM proteins, play important roles in shaping epithelia, especially narrow epithelial tubes (Sundaram and Cohen, 2017). This led us to consider whether epithelial features of amphid neurons and glia might be important for their development.

Here, we show that amphid neurons and glia exhibit hallmarks of epithelial cells, including tight junctions and apical-basal polarity. Consistent with the idea that DYF-7 acts as a component of apical ECM, we find that it forms fibrils at the apical surface of the developing amphid. Loss of DYF-7 leads to rupture of the tube-shaped glial pore that surrounds the dendrites, reminiscent of rupture defects seen in other epithelial tubes upon disrupting apical ECM. These defects are exacerbated by loss of FRM-2, a homolog of proteins that are also involved in epithelial integrity in other systems. Finally, we show that a requirement for DYF-7 is shared by other sensory neurons that are exposed to the environment. Our results suggest that these environmentally-exposed sensory neurons and their associated glia should be viewed as part of an epithelium that is continuous with the skin, with apical-basal polarity and a mechanism of development that shares key features with other epithelia, especially narrow epithelial tubes.

## RESULTS

### Amphid neurons and glia exhibit hallmarks of an epithelium

In the mature amphid, dendrites are exposed to the environment through a tube-shaped glial pore (Fig. 1A-D). We asked whether amphid neurons and glia exhibit properties of true epithelial cells, namely the presence of tight junctions and apical-basal polarity.

**Figure 1.**
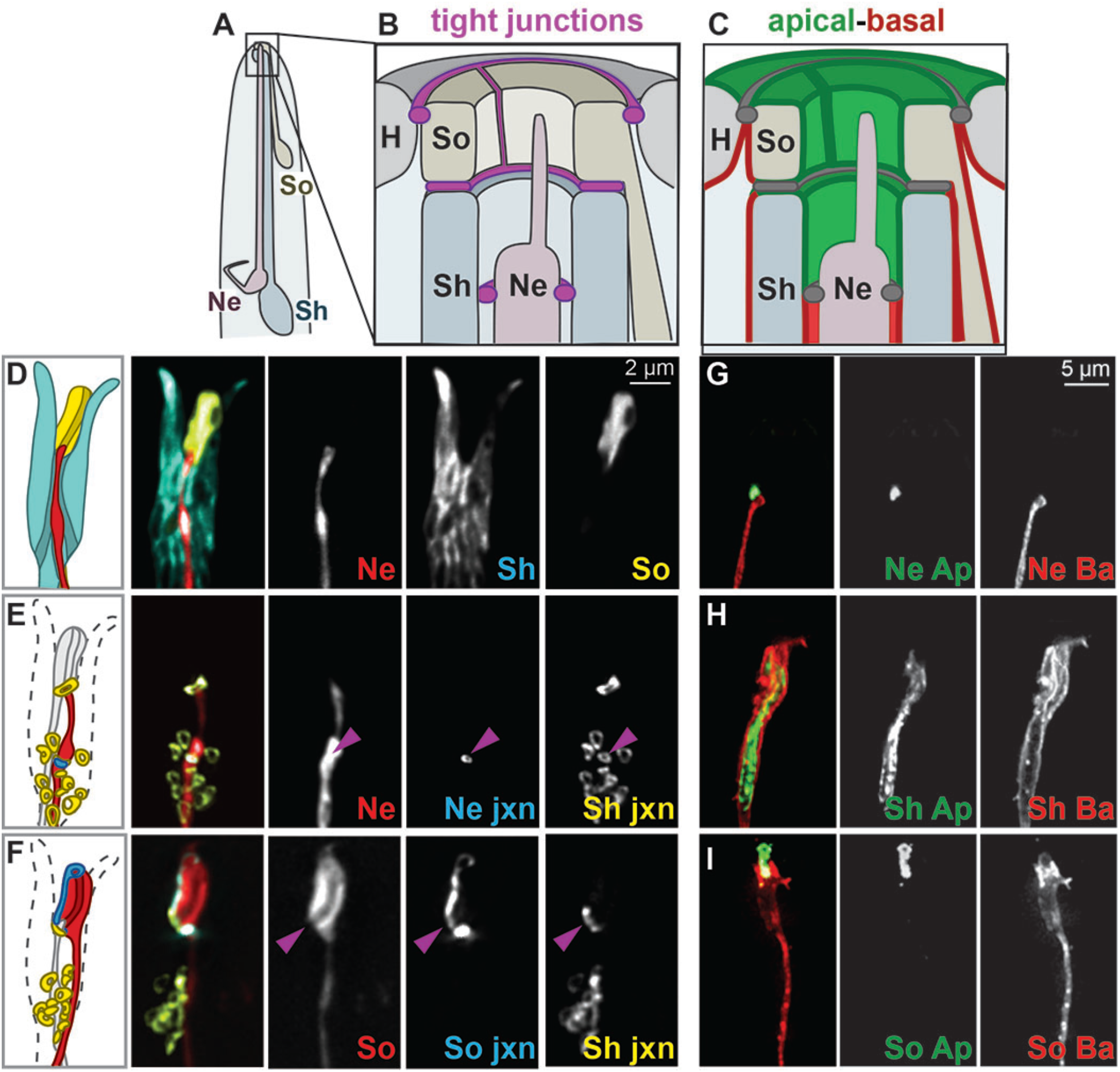
Amphid neurons and glia exhibit tight junctions and distinct apical and basolateral surfaces. (A-C) Schematics showing relative positions of (A) amphid dendrite endings, (B) tight junctions (purple), and (C) apical (green) and basolateral (red) surfaces. For simplicity, only a single neuron is drawn. (D) Cell-specific markers were used to visualize single neurons and glia. ASER neuron (*gcy-5*pro, red); sheath glia (*F16F9.3*pro, blue); socket glia (*grl-2*pro, yellow). (E,F) Overlap of neuron:sheath and sheath:socket tight junctions was visualized by using these promoters to express AJM-1-YFP in the sheath and cytoplasmic mCherry plus AJM-1-CFP in (E) the neuron or (F) the socket. Dotted line in schematics, approximate outline of sheath; purple arrowheads, position of overlapping tight junctions. (G-I) The apical and basolateral markers ApiGreen (Ap, green) and BasoRed (Ba, red) (see Methods and Supp. Fig. S1D) were expressed in (G) the neuron, (H) the sheath, (I) the socket using the same promoters. Ne, neuron; Sh, sheath glia; So, socket glia; H, hypodermis (skin); jxn, junction; Ap, ApiGreen; Ba, BasoRed.

#### Tight junctions

Classical EM analysis described electron-dense tight junctions between amphid neurons and the sheath glial cell; the sheath and socket glia; and the socket and skin (Ward et al., 1975; Perkins et al., 1986). Consistent with these observations, recent studies noted localization of the tight junction protein AJM-1 near amphid dendrite endings (Nguyen et al., 2014; Nechipurenko et al., 2016). We used promoters specific to the ASE neuron, the sheath, or the socket (*gcy-5*pro, *F16F9.3*pro, and *grl-2*pro, respectively; Fig. 1D) to drive expression of a portion of AJM-1 cDNA tagged with CFP or YFP in each of these cell types. This approach allowed us to visualize defined tight junctions with single-cell resolution.

AJM-1-YFP expressed in the sheath glial cell labeled 13 ring-shaped junctions (Fig. 1E, yellow). Their positions suggest a single large junction to the socket plus 12 smaller junctions to each of the dendrites. The ASE dendrite passes through one of these smaller rings (Fig. 1E, red). AJM-1-CFP expressed in the ASE neuron labels a ring that is concentric with one of the sheath rings and is positioned near the dendrite ending, proximal to the cilium (Fig. 1E, arrowhead). We obtained similar results with other amphid neurons (AWB and AWC; Supp. Fig. S1A). These results indicate that each neuron forms an individual ring-shaped tight junction with the sheath glial cell.

AJM-1-CFP expressed in the socket glial cell labels two ring-shaped structures whose positions are consistent with the presence of socket:skin and socket:sheath junctions (Fig. 1F, blue). The latter overlaps AJM-1-YFP expressed in the sheath glial cell (Fig. 1F, yellow). AJM-1-CFP also labels a stripe connecting these rings, consistent with the auto-junction that the socket makes on itself as it wraps into a single-cell tube (Fig. 1B,F). We obtained similar results using another tight junction component (DLG-1, Supp. Fig. S1B). Together, the neuron:sheath, sheath:socket, and socket:skin tight junctions delineate an outward-facing surface continuous with the skin epithelium (Fig. 1B).

#### Apical-basal polarity

Another hallmark of epithelia is the presence of biochemically distinct apical and basolateral membranes. We recently developed constructs derived from the transmembrane protein SAX-7 that localize exclusively apically or basolaterally in epithelia (Supp. Fig. S1D). For simplicity, we will refer to these markers as ApiGreen and BasoRed. We expressed these markers under control of promoters specific to single amphid neurons or glia.

In the neuron, ApiGreen localized distal to the tight junction in the dendrite ending (Fig. 1G, green). BasoRed localized along the length of the dendrite and was excluded from the dendrite ending (Fig. 1G, red). This suggests that the outward- and inward-facing surfaces of the neuron are biochemically distinct. The region labeled by ApiGreen has been referred to as the periciliary membrane compartment (PCMC) and is a site of membrane trafficking important for ciliogenesis (Blacque and Sanders, 2014). ApiGreen was typically, but not always, excluded from the cilium; in general, transmembrane proteins are excluded from the cilium via a diffusion barrier called the ciliary gate (Garcia-Gonzalo and Reiter, 2017) but higher expression levels of ApiGreen may have allowed it to leak through this barrier in some cases.

In the sheath glial cell, ApiGreen localized to the outward-facing surface between the sheath:neuron and sheath:socket junctions (Fig. 1H, green). A marker of apical cytoskeleton, ERM-1, localized in the same pattern (Supp. Fig. S1C). BasoRed localized to the rest of the glial surface and did not overlap with ApiGreen (Fig. 1H, red). Similarly, in the socket glial cell, ApiGreen localized to the outward-facing surface between the sheath:socket and socket:skin junctions, with BasoRed localizing to the rest of the glial surface (Fig. 1C,I).

Together, the presence of tight junctions and apical-basal polarity suggest that amphid neurons and glia can be viewed as part of an epithelium continuous with the skin.

### DYF-7 is an apical ECM component that prevents rupture of the amphid epithelium

Next, we considered how the epithelial properties of these neurons and glia might relate to their morphogenesis in the embryo. We previously found that the ZP domain protein DYF-7 is required for amphid dendrite extension during development. Most ZP domain proteins are found at the apical surfaces of epithelia, where they polymerize into fibrils and play important roles in epithelial morphogenesis, including preventing rupture of narrow epithelial tubes (Plaza et al., 2010). Viewing the amphid neurons and glia as part of an epithelium, we tested three predictions about DYF-7: first, that it localizes to apical surfaces; second, that it polymerizes into fibrils; and third, that defects caused by loss of DYF-7 reflect epithelial rupture.

#### DYF-7 localizes to apical surfaces

We examined the localization of DYF-7 in the developing amphid and other tissues. Most ZP domain proteins are synthesized with a membrane anchor and undergo proteolytic cleavage at a consensus furin cleavage site (CFCS) to release the ectodomain (Bokhove and Jovine, 2018). DYF-7 contains a CFCS and can undergo proteolytic cleavage *in vitro* (Heiman and Shaham, 2009). In order to determine whether DYF-7 undergoes cleavage *in vivo* and to track the localization of its ectodomain, we tagged DYF-7 with superfolderGFP (sfGFP) on its ectodomain (DYF-7^ecto^) and mCherry on its cytoplasmic tail (DYF-7^cyt^) (Fig. 2A). This construct completely rescues a *dyf-7* null mutant (Supp. Fig. S2A).

**Figure 2.**
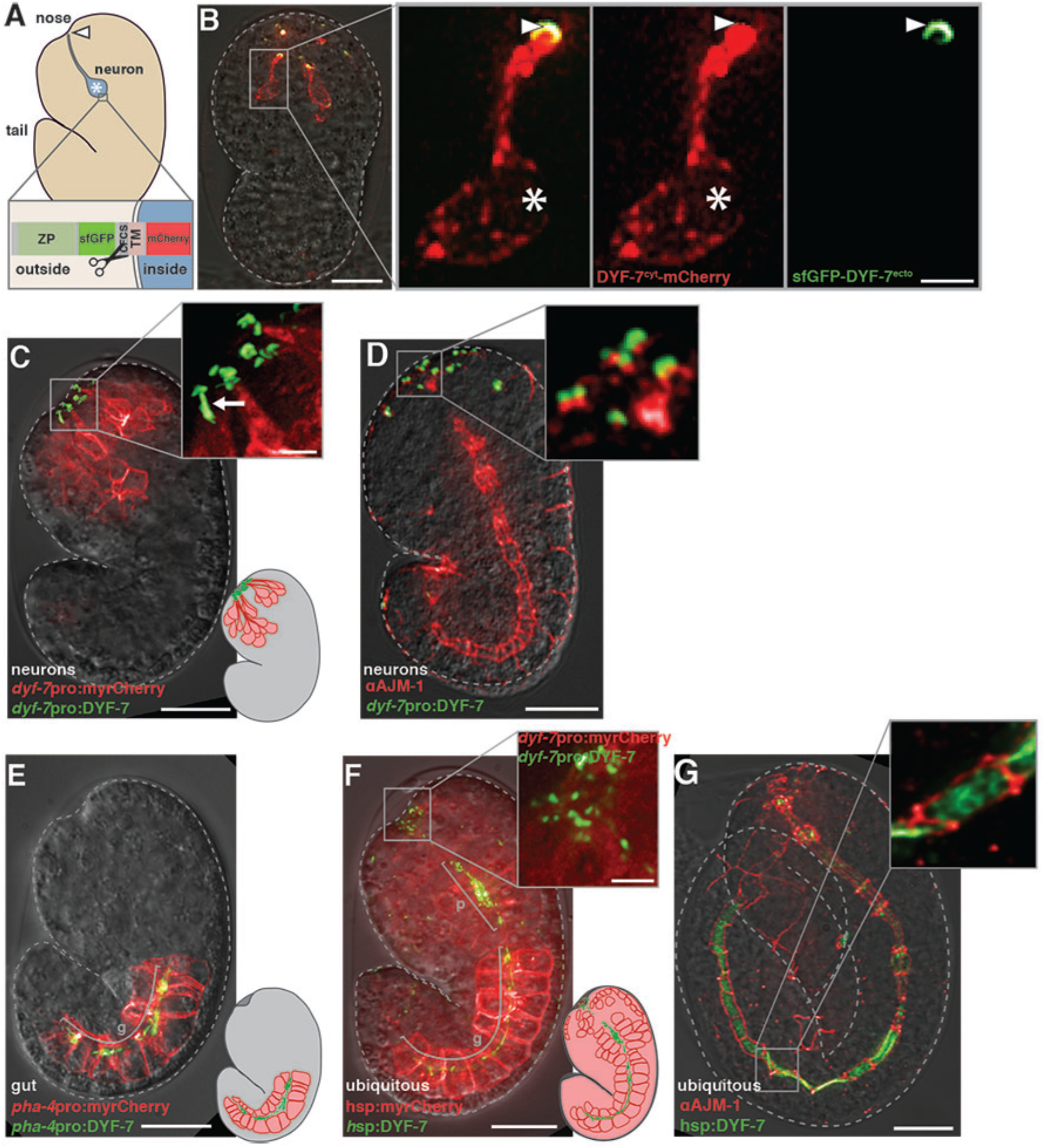
DYF-7 is cleaved *in vivo* and its ectodomain localizes to apical surfaces. (A) Cartoon schematic of an embryo showing a single neuron (blue; cell body, *) with its dendrite extending to the developing nose (dendrite ending, arrowhead). Topology of the DYF-7 reporter construct, consisting of the ZP domain (ZP), superfolder GFP tag (sfGFP), consensus furin-family cleavage site (CFCS), transmembrane segment (TM), and cytoplasmic region tagged with mCherry (mCherry). (B) A live, intact embryo expressing this construct in sensory neurons (*dyf-7*pro) during dendrite extension. (C,D) Embryos expressing sfGFP-DYF-7 in sensory neurons (*dyf-7*pro) (C) with myristyl-mCherry or (D) with immunostaining against AJM-1. Some structures appear as short linear segments (arrow) rather than rounded caps. (EG) Embryos misexpressing sfGFP-DYF-7 in (E) gut (*pha-4*pro) with myristyl-mCherry or (F,G) ubiquitously (heat shock promoter) (F) with myristyl-mCherry or (G) with immunostaining against AJM-1. Brackets mark pharynx (p) and gut (g). Scale bars: 10μm, main panel; magnified insets.

We expressed this construct under control of the *dyf-7* promoter, which is expressed in sensory neurons, and examined embryos at the time of dendrite extension. We found that the DYF-7^ecto^ and DYF-7^cyt^ fragments localized differently, consistent with proteolytic cleavage at the CFCS (Fig. 2B). Deletion of the CFCS led to increased colocalization, confirming that DYF-7 undergoes CFCS-dependent proteolysis *in vivo* (Supp. Fig. S2B). The membrane-anchored cytoplasmic tail of DYF-7 localized diffusely across the neuronal membrane with enrichment at dendrite endings (Fig. 2B, red), consistent with our previous observations (Heiman and Shaham, 2009). In contrast, the ZP domain of DYF-7^ecto^ localized with exquisite precision to caps at dendrite endings (Fig. 2B, green). This localization requires the ZP domain, as a construct lacking the ZP domain localized diffusely throughout the extra-embryonic space (Supp. Fig. S2C).

Multiple dendrite caps were observed, corresponding to the amphid and other sense organs in the head (Fig. 2C). In some cases, these structures took the shape of short linear segments, consistent with the luminal surface of a tube (Fig. 2C, arrow). Immunostaining with an antibody against AJM-1 revealed that each of these caps is situated adjacent to a tight junction (Fig. 2D). The position of these caps thus corresponds to the outward-facing apical surfaces of the dendrites.

Next, we asked where DYF-7 would localize if it were misexpressed in other epithelia. We expressed the tagged DYF-7^ecto^ construct under control of gut-specific or ubiquitously-expressed promoters. In the gut, DYF-7^ecto^ localized to the lumen, which is the apical surface of this well-studied model epithelium (Leung et al., 1999) (Fig. 2E). When expressed ubiquitously, DYF-7^ecto^ localized to the lumens of the gut and pharynx as well as to dendritic caps at the developing nose (Fig. 2F). It was absent from non-epithelial regions of the embryo. Immunostaining against AJM-1 confirmed that misexpressed DYF-7^ecto^ localizes in the pharynx and gut specifically to apical surfaces (Fig. 2G).

#### *DYF-7 forms fibrils* in vivo *and* in vitro

Other ZP domain proteins contribute to apical ECM by polymerizing into fibrils that form a meshwork at the apical surfaces of epithelia (Jovine et al., 2002; Jovine et al., 2004; Jovine et al., 2006; Schaeffer et al., 2009). We therefore asked whether DYF-7 also forms fibrils.

Previous EM studies noted the appearance of extracellular fibrils in the developing amphid (Oikonomou et al., 2011). However, these structures are difficult to visualize using standard approaches because their appearance depends on the angle of sectioning and the developmental stage of the embryo. We used a method called positional correlative anatomy EM to provide greater control over these variables (Kolotuev, 2014; Burel et al., 2018). Briefly, a high-pressure frozen sample is flat-embedded at the surface of a resin block, a region of interest is identified at high magnification, and the sample is then oriented for ultramicrotome sectioning with respect to known anatomical features.

Using this method, we identified embryos undergoing dendrite extension — between the comma and 1.5-fold stages of embryogenesis (~420 min post-fertilization) — and sectioned them at an angle parallel to the long axis of the dendrites (Fig. 3A, top). In wild-type animals at this stage, cilia had not yet formed but a centriole/basal body was always present at the dendrite endings (annotated in Fig. 3Aiii, arrowheads). We found that the developing dendrites form direct contacts on each other, rather than making individual junctions to the sheath as in the mature structure. The dendrites are bundled together in a single channel of the sheath glial cell that is continuous with the channel of the socket glial cell, which in turn is open to the extraembryonic space (Fig. 3A). In all cases, we observed fibrils that originate near the dendrite endings, extend through the sheath:socket channel, and terminate near the extra-embryonic ECM, extending ~1.5μm in total (Fig. 3A; fibrils, brackets). These fibrils were consistently visible across serial sections in all embryos imaged (n=3 amphids).

**Figure 3.**
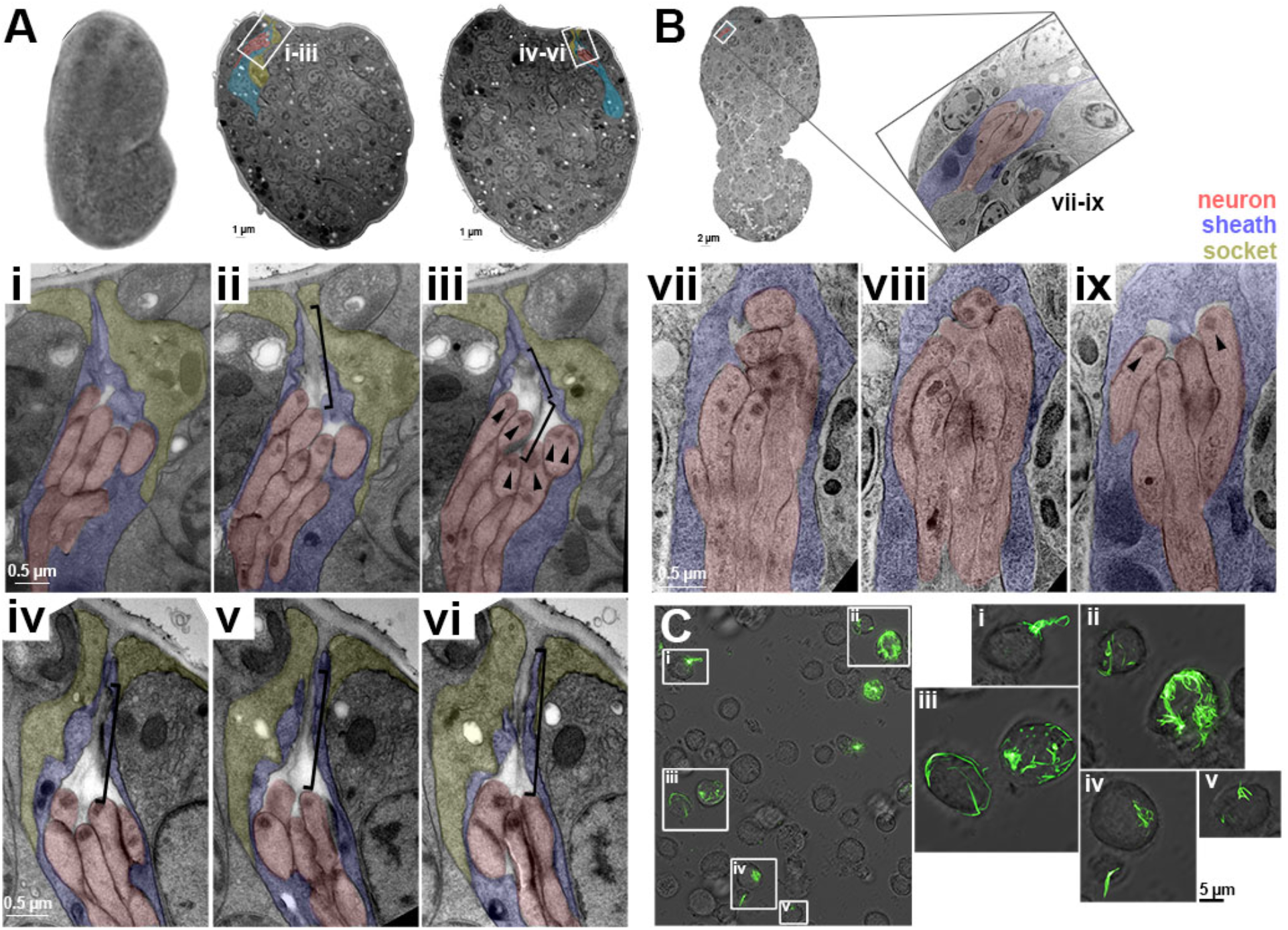
DYF-7 forms extracellular fibrils. (A) Image of a wild-type embryo after high-pressure freezing, fixation, and flat embedding. Flat embedding allowed selection of embryos at the desired stage and orientation for sectioning. Serial sectioning led to identification of the (i-iii) left- and (iv-vi) right-hand amphids. Three serial sections of each amphid are shown. Neurons, red; sheath, blue; socket, yellow. Basal bodies/centrioles are annotated in (iii) with arrowheads. Extracellular fibrils are marked with brackets. (B) Three serial sections (viii-ix) of an amphid in a *dyf-7* embryo at the same developmental stage. The sheath forms a pocket around the neurons and no fibrils are visible. Basal bodies/centrioles appear normal (arrowheads). (C) *Drosophila* S2 cells were transfected with sfGFP-DYF-7. A representative field of cells is shown. GFP-labeled structures appeared as (i) a plume at the cell surface, (ii) complex tangles, (iii) puncta and fibrils that appear to encircle the cells, (iv, v) small webs.

Next, we compared this to *dyf-7* mutant embryos at the same stage. In *dyf-7* embryos at this stage, the nascent dendrites and sheath have just begun to withdraw from the developing nose (Fig. 3B, top). The dendrites are organized within the sheath in a manner similar to wild type but the sheath forms a closed cavity rather than an open channel (Fig. 3Bvii-ix). We did not observe fibrils in any of the *dyf-7* embryos imaged (n=6 amphids). We considered the possibility that fibrils might be present but disorganized and hard to recognize (e.g., too short, or oriented
randomly). However, we were unable to identify fibril-like structures in any serial sections across a variety of embryo orientations and a range of section thicknesses (60-100nm). This finding suggests that DYF-7 is required for formation of these fibrils.

Finally, we asked whether DYF-7 is sufficient to form fibrils *in vitro*. We expressed sfGFP-DYF-7 in *Drosophila* S2 cells. We observed GFP-positive clumps and tangles extending from the surfaces of cells (Fig. 3C). In some cases, GFP-positive fibrils appeared to wrap around the cells (Fig. 3Cii, iii). Thus, our results indicate that DYF-7 is sufficient to form fibrils *in vitro*, and that it is necessary for the formation or maintenance of such fibrils at the apical surface of the developing amphid *in vivo*. These observations support the idea that, like ZP domain proteins in other systems, DYF-7 is an apical ECM component.

#### DYF-7 is required to prevent rupture of the tube-shaped glial pore

Studies in *C. elegans* and *Drosophila* have shown that disrupting ZP domain proteins that are part of the apical ECM leads to epithelial rupture, including loss of a continuous lumen in the narrow epithelial tubes of the *C. elegans* excretory system and the *Drosophila* tracheal system (Wilkin et al., 2000; Jaźwińska et al., 2003; Bökel et al., 2005; Kelley et al., 2015; Gill et al., 2016). Previously, we focused on the role of DYF-7 in anchoring dendrites to the embryonic nose, however, viewing DYF-7 as an apical ECM component, we decided to ask if dendrite anchoring defects might be related to rupture of the tube-shaped glial pore that normally surrounds the dendrites.

In *dyf-7* mutants, amphid dendrites and the sheath glial cell fail to extend but the socket glial cell still extends a process to the nose (Heiman and Shaham, 2009) (Fig. 4A). First, we examined tight junctions and apical-basal markers in the socket. At the nose, tight junctions were still present between the socket and the skin, and the apical marker ApiGreen still localized to the outward-facing surface (Fig. 4B,C). However, the auto-junction that normally seals the socket into a tube was absent and no signs of the normal socket channel could be seen. In 32 of 48 amphids, the sheath and socket appeared completely dissociated from each other; however, in the remaining 16 amphids, the socket glial cell extended an ectopic posterior process to contact the shortened sheath glial cell (see schematic in Fig. 4A). In these examples, tight junction markers colocalized at the sheath:socket contact, but appeared as a small dot rather than a ring as in wild type (Fig. 4D). ApiGreen did not localize to the sheath:socket contact (Fig. 4E). These results suggest that the socket cell retains tight junctions and apical-basal polarity, but no longer forms a continuous channel with the sheath.

**Figure 4.**
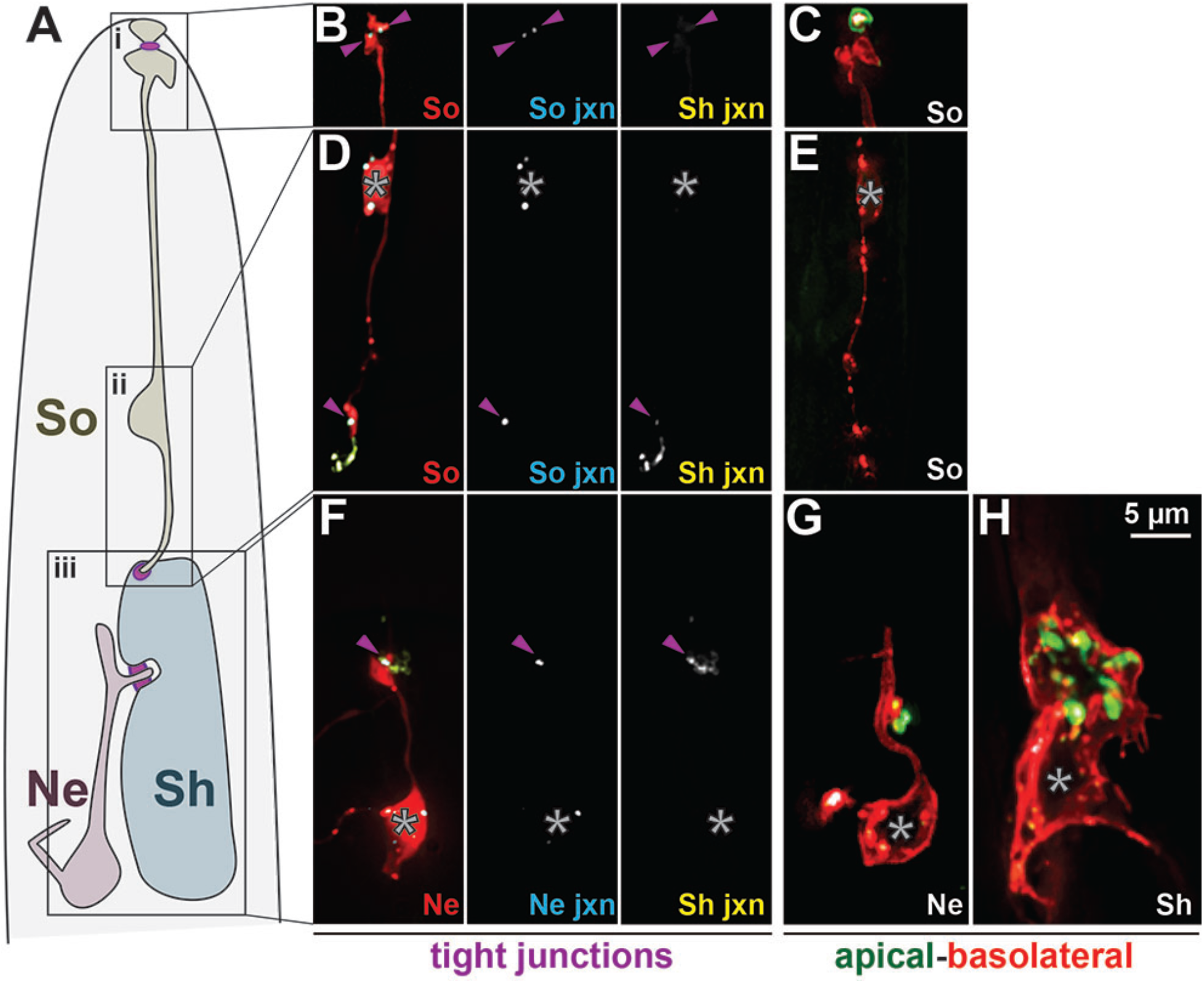
Disruption of the amphid glial channel in *dyf-7* mutants. (A) Schematic showing the relative positions of the socket (So), sheath (Sh), and neurons (Ne) in a *dyf-7* mutant. Posterior process from socket to sheath occurred in 16/48 amphids. Regions of interest are boxed: (i) socket:skin junction, shown in (B,C); (ii) sheath:socket junction, shown in (D,E); (iii) neuron:sheath junction, shown in (F-H). (B,D,F) AJM-1-YFP expressed in sheath plus cytoplasmic mCherry and AJM-1-CFP expressed in (B,D) socket or (F) neuron. (C,E,G,H) ApiGreen and BasoRed expressed in (C,E) socket, (G) neuron, or (H) sheath. All markers and promoters are the same as in Fig. 1. Purple arrowheads, positions of relevant tight junctions; asterisks, cell bodies.

Next, we examined these markers in the amphid sheath and an amphid neuron. Both cells retained their tight junctions and apical-basal polarity (Fig. 4F-H). The dendrite still appeared to enter the sheath and form tight junctions with it, consistent with our previous EM analysis (Heiman and Shaham, 2009) (Fig. 4F). However, the apical surface of the sheath glial cell appeared to be organized in a network of cyst-like internal pouches rather than an open channel (Fig. 4H). These defects are reminiscent of the loss of lumen integrity caused by disruption of ZP domain proteins in other narrow epithelial tubes.

Together, these results strongly suggest that amphid neurons and glia use a developmental mechanism that shares key features with other epithelia, namely, the use of a fibril-forming ZP domain protein in the apical ECM to prevent rupture of a narrow epithelial tube.

### DYF-7 defects are enhanced by loss of FRM-2, a homolog of epithelial integrity proteins

Next, we wanted to identify additional factors that help to promote morphogenesis of amphid neurons and glia. We reasoned that, if the amphid shares developmental features with epithelia, then it might use additional factors that are typically found in developing epithelia, such as proteins involved in apical-basal polarity or cell junction integrity. Conversely, if the amphid differs in important ways from conventional epithelia, then it might rely more heavily on factors that are typically associated with neuronal or glial development, or novel factors.

To this end, we performed an unbiased genetic screen using a sensitized background in which DYF-7 activity is mildly impaired. We took advantage of a hypomorphic missense allele, *dyf-7(ns117)*, in which only ~35% of amphids are affected (Heiman and Shaham, 2009). Briefly, we performed chemical mutagenesis of *dyf-7(ns117)* mutants bearing a fluorescent marker for a single amphid neuron. 1000 F2 self-progeny bearing novel unique homozygous mutations were picked to individual plates and their clonal F3 offspring were visually screened for increased (enhanced) penetrance of the dendrite extension defect (Fig. 5A). We isolated two mutants with heritably increased penetrance: *hmn162* and *hmn169*. As *hmn169* was less strongly enhanced and exhibited semi-dominance (see Methods), we chose to focus on *hmn162*.

**Figure 5.**
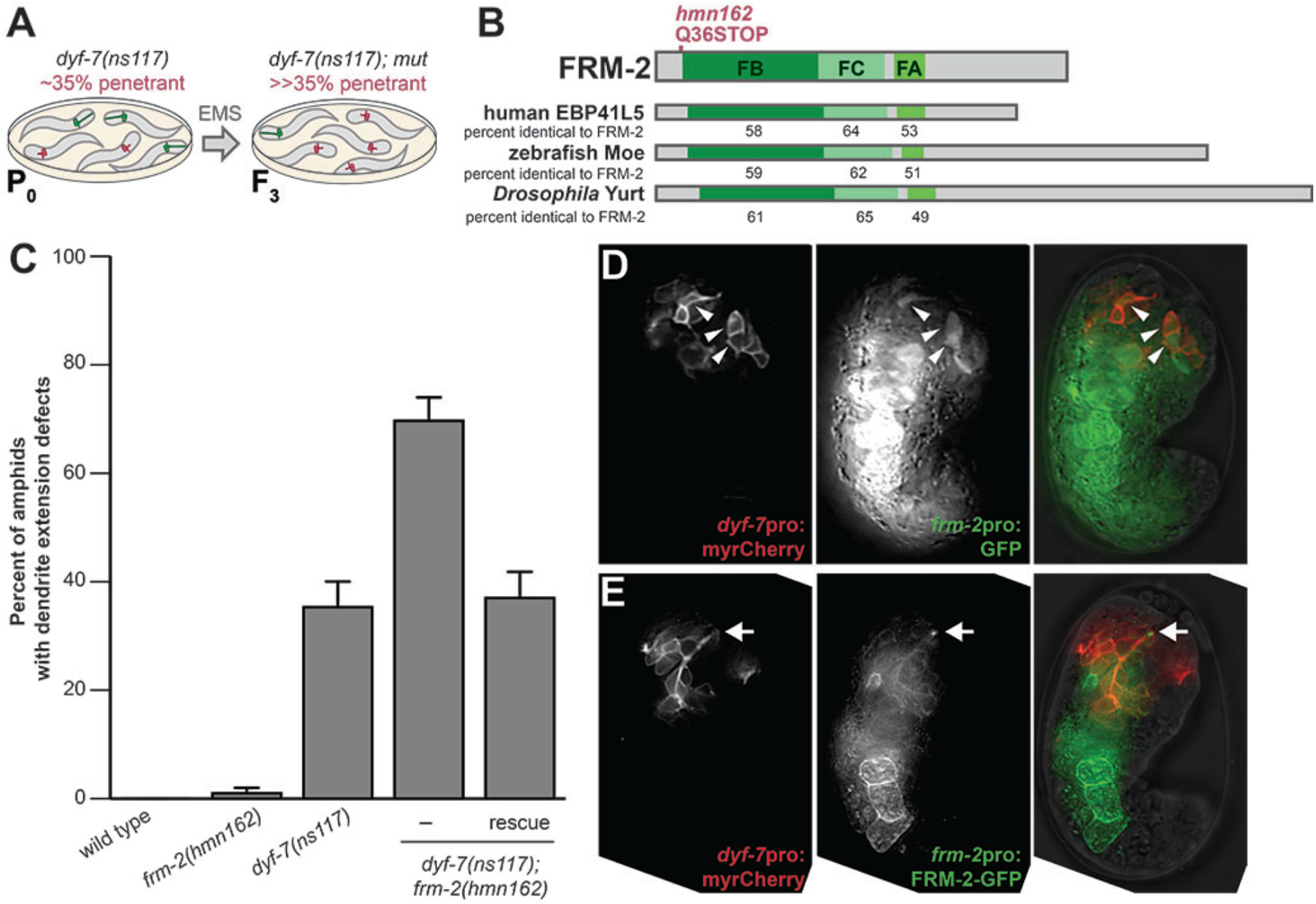
Loss of FRM-2 enhances weak *dyf-7* defects. (A) Schematic of genetic strategy used to isolate enhancers of the hypomorphic missense mutant *dyf-7(ns117)*. Parental (P0) strain exhibits ~35% short amphids (red) and ~65% full-length amphids (green). Following mutagenesis, clonal third generation (F3) progeny bearing additional mutations (mut) were visually screened for enhanced penetrance. (B) Schematic of the FRM-2 protein showing location of *hmn162* nonsense mutation. FRM-2 is roughly 50-60% identical to human EPB41L5, zebrafish moe, and *Drosophila* Yurt throughout the FERM B-lobe (FB), FERM C-lobe (FC), and FERM-adjacent (FA) domains, with a more divergent carboxyterminal sequence. (C) *frm-2(hmn162)* enhances the amphid dendrite extension defects of *dyf-7(ns117)*. “rescue”, animals bearing a transgene with a wild-type *frm-2* genomic fragment. n≥100 amphids per genotype. Error bars, standard error of the mean. (D) *frm-2*pro drives expression in sensory neurons (*dyf-7*pro, arrowheads) at the time of dendrite extension, as well as bright expression in pharynx and gut. (E) A rescuing FRM-2-GFP construct expressed by its endogenous promoter localizes to developing dendrite endings (arrow).

Single-step mapping and sequencing of *hmn162* identified a nonsense mutation in a previously uncharacterized gene called *frm-2* (Fig. 5B). Mutants bearing *frm-2(hmn162)* alone exhibit very rare (1±1%) dendrite defects (Fig. 5C). However, whereas *dyf-7(ns117)* exhibits 35±5% penetrant dendrite defects, the *dyf-7(ns117); frm-2(hmn162)* double mutant exhibits an enhanced penetrance of 70±4% (Fig. 5C). A wild-type *frm-2* genomic fragment expressed under its own regulatory sequences completely rescues this phenotype, returning the penetrance to the level of the *dyf-7(ns117)* single mutant (37±5%) (Fig. 5C).

*frm-2* is predicted to encode a FERM-domain protein most closely related to mammalian EPB41L5, zebrafish mosaic eyes (moe), and *Drosophila* Yurt (Yrt) (Fig. 5B). Each of these proteins has been shown to interact with the Crumbs apical polarity complex to regulate epithelial morphogenesis (Hsu et al., 2006; Laprise et al., 2006; Gosens et al., 2007; Laprise et al., 2009; Salis et al., 2017). Disruption of EPB41L5, moe, and Yrt has been shown in each of these systems to cause disorganized tight junctions and loss of epithelial integrity (Jensen and Westerfield, 2004; Salis et al., 2017; Schell et al., 2017). Interestingly, in *Drosophila* trachea, the relationship between Yrt and apical ECM has been examined, and these pathways were found to act independently and in parallel to shape epithelial tubes, an observation that would be consistent with the loss of FRM-2 enhancing *dyf-7* defects (Laprise et al., 2010).

To determine whether FRM-2 might be acting directly in the amphid, we examined its expression. At the time of dendrite extension, a *frm-2*pro:GFP reporter is expressed brightly in the developing epithelium of the pharynx and gut, with much weaker but consistent expression in sensory neurons (Fig. 5D, arrowheads). We examined FRM-2 protein localization using a *frm-2*pro:FRM-2-GFP reporter. We consistently observed localization at amphid dendrite endings (Fig. 5E, arrow), although it was sometimes obscured by the much brighter signal from the pharynx and gut.

These results support the idea that FRM-2 acts apically in the developing amphid to prevent epithelial rupture, similar to what has been shown for EPB41L5, moe, and Yrt in other epithelia. Our results are consistent with the idea that, like Yrt, FRM-2 acts in parallel to apical ECM proteins like DYF-7 to regulate the formation of a narrow epithelial tube. While our unbiased screen was not comprehensive, it is interesting that it identified a factor that most closely resembles ones found in other epithelia, rather than ones typically involved in the development of neurons or glia.

### Other environmentally-exposed neurons also require DYF-7

In addition to the amphid, other sensory neurons in the head are exposed directly to the environment and are thus likely to have epithelial properties (Ward et al., 1975). These include the cephalic (CEP), inner labial (IL), and outer labial (OL) neurons. Each of these classes of neuron has its own dedicated glia (CEP sheath and socket; IL sheath and socket; OL sheath and socket) that form a tube-shaped channel around the dendrites. In all cases, there are junctions between the dendrites and sheath; the sheath and socket; and the socket and skin to delineate an outward-facing surface (Ward et al., 1975). We wondered if these neurons also require DYF-7.

To test this idea, we used cell-specific promoters to visualize amphid, CEP, OL, and IL dendrites in wild-type and *dyf-7* animals (Fig. 6). In *dyf-7* mutants, amphid neurons are typically ~10% of wild-type length (Fig. 6B,C). We found that CEP, OL, and IL neurons are also affected by the loss of DYF-7 (Fig. 6B,C). The defects were highly penetrant, with >90% of dendrites having lengths at least five standard deviations below the wild-type mean (Fig. 6C). These dendrites were typically ~80% of wild-type length (Fig. 6C), but length varied even within a class; for example, ventral CEPs were consistently more affected than dorsal CEPs (75.2±17.4 and 91.2±3.2 percent of nose length, respectively).

**Figure 6.**
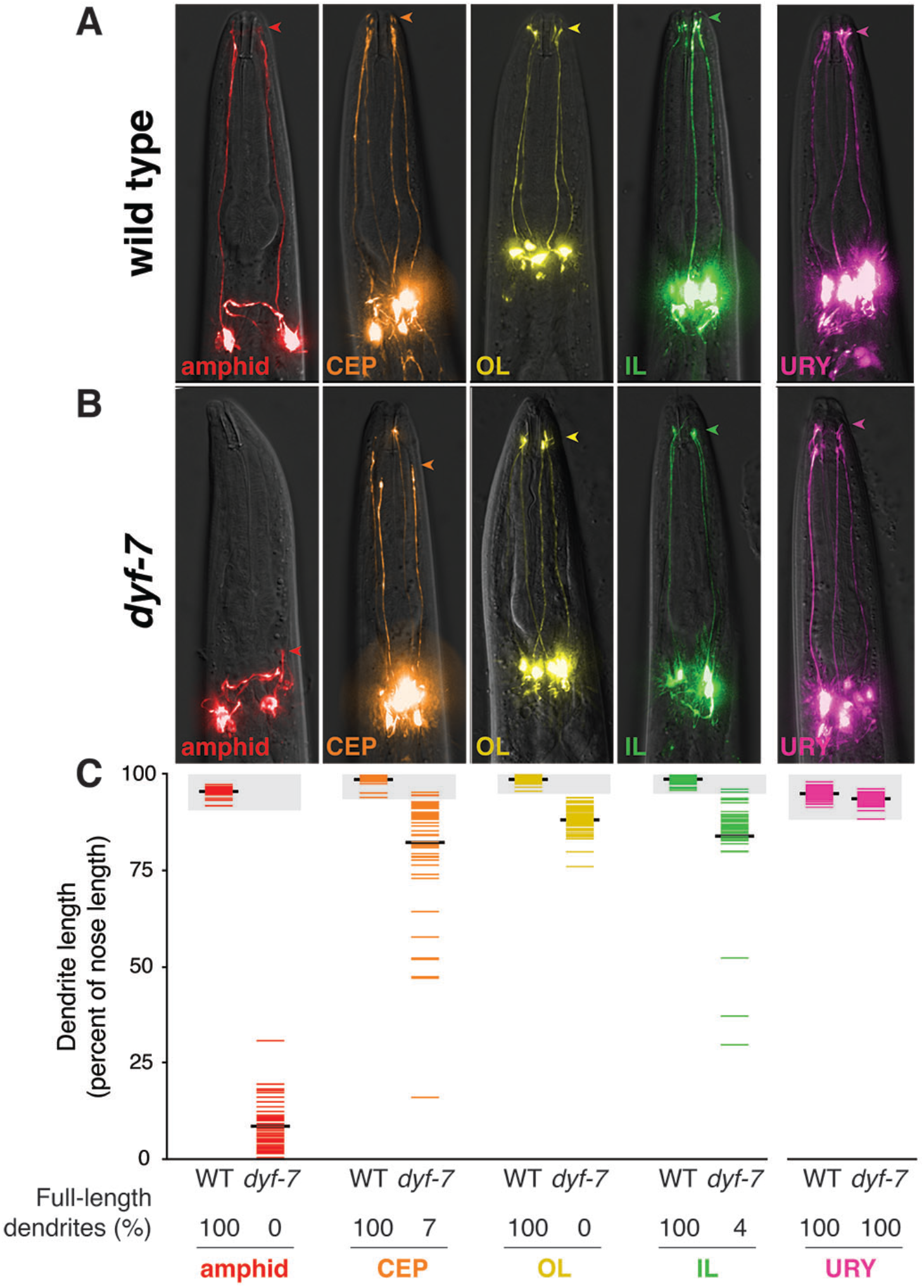
Sensory neurons within epithelia exhibit a shared dependence on DYF-7. Cell-specific markers were used to visualize sensory neurons within epithelia – amphid (AWC, *odr-1*pro), CEP (*dat-1*pro), OL (OLQ, *ocr-4*pro), and IL (IL2, *klp-6*pro) – as well as URY (*tol*-*1*pro), which is not within an epithelium, in (A) wild-type and (B) *dyf-7(ns119)* null mutants. Arrowheads, position of dendrite endings. (C) Quantification of dendrite lengths as a fraction of nose length for each neuron in wild-type (left column) and *dyf-7* (right column) animals. Colored bars represent individual dendrites (n≥48 per column); black bars represent means. Shaded areas represent the wild-type mean ± 5 standard deviations (SD) for each neuron type; the percent of dendrites in this range (“full-length dendrites”) is given.

For comparison, we also examined the URY sensory neuron. This neuron extends an unbranched dendrite to the nose that associates with sheath glia, but its dendrite ending does not enter the glial channel and is not exposed to the environment; thus, it is not expected to have epithelial properties (Ward et al., 1975). We found that these dendrites were mostly unaffected by loss of DYF-7, suggesting that its role is specific to environmentally-exposed neurons (Fig. 6B,C).

Together, these results indicate that the amphid, CEP, OL, and IL neurons share a dependence on DYF-7. Thus, what we have learned about epithelial features of amphid neurons and glia might apply broadly to other environmentally-exposed neurons in *C. elegans* and, perhaps, in other organisms as well.

## DISCUSSION

### Epithelial properties of amphid neurons and glia

Our results suggest that amphid neurons and glia exhibit features of an epithelium, including tight junctions and apical-basal polarity. Similar anatomical arrangements are found in the olfactory and vomeronasal systems of mammals and are the predominant type of sensory structure in invertebrates, including the campaniform, trichoid, and chordotonal neurons of *Drosophila* and most sensory neurons of *C. elegans* (IL, OL, CEP in the head as discussed above, as well as phasmids, deirids and male-specific tail sensory neurons) (White et al., 1986; Hartenstein, 1988). Thus, neurons with epithelial properties may be an ancestral type of sensory structure.

Viewed as an epithelium, the glial pore of the amphid especially resembles narrow epithelial tubes found in capillaries and the branching ductwork of kidney, pancreas, lung, or other organs (Sundaram and Cohen, 2017). The development of narrow epithelial tubes has been studied extensively in the *C. elegans* excretory system, which includes three tube-forming cells: the canal, duct, and pore cells (Sundaram and Buechner, 2016). The canal and pore cells bear anatomical similarities to the amphid sheath and socket, respectively, with the canal and sheath forming seamless tubes that are not lined with cuticle, and the pore and socket using auto junctions to create cuticle-lined tubes that are open to the environment via junctions with the skin. An additional curious parallel is that the excretory pore and amphid socket can each act as stem cells, dividing post-embryonically to give rise to neurons (Sulston and Horvitz, 1977; Sammut et al., 2015). Besides their anatomical similarities, the excretory system and amphid share genetic and molecular features: Several mutants have been described that cause strong phenotypes affecting both the excretory system and amphid (Rdy phenotype, for “rod-like larval lethality and dye-filling defective”), and several molecular markers exhibit prominent expression shared by both these organs (e.g., *pros-1*, *daf-6*, *lin-48, grl-2*) (Michaux et al., 2000; Johnson et al., 2001; Perens and Shaham, 2005; Hao et al., 2006b; Liégeois et al., 2007; Kolotuev et al., 2013; Kage-Nakadai et al., 2016; Wallace et al., 2016).

These parallels are consistent with the idea that the development of the amphid resembles that of other narrow epithelial tubes. With its easily visualized morphology and the ability to label or genetically manipulate each neuron and glial cell with single-cell specificity, the amphid offers a novel and potentially powerful model system with which to ask questions about epithelial development. The epithelial properties of amphid neurons also raise new questions, including (1) how can a cell simultaneously have apical-basal and axon-dendrite polarity?; (2) given that the transcription factor LIN-26 is thought to promote epithelial fates while repressing neuronal fates(Labouesse et al., 1994; Labouesse et al., 1996), what kind of transcriptional programs allow the simultaneous expression of epithelial and neuronal genes?; and (3) why do DYF-7 and FRM-2 affect epithelia that contain neurons and glia more strongly than they affect other epithelia?

### DYF-7 as an apical ECM component

ZP domain proteins are found in the apical ECM of nearly all epithelia and are present from cnidaria to chordates (Plaza et al., 2010; Matveev et al., 2012). They play important roles in human disease: the tectorins of the inner ear are mutated in deafness; the tumor suppressor DMBT1 is expressed in many epithelia and is frequently mutated in cancer; uromodulin in the kidney is mutated in hereditary nephropathy; GP-2 is expressed in the pancreas and intestine and acts as an autoantigen in Crohn’s disease; and the TGF-β co-receptor endoglin is expressed in endothelia and is mutated in vascular dysplasia and elevated in pre-eclampsia (Legan et al., 1997; Ligtenberg et al., 2007; Gregory et al., 2014; Roggenbuck et al., 2014; Devuyst et al., 2017). Previously, it was unclear how the expression of DYF-7 in neurons fit with this well-established literature on ZP domain proteins in the apical ECM of epithelia. Our results clarify that, like other ZP domain proteins, DYF-7 localizes to apical surfaces and polymerizes into fibrils, and thus it is likely to be an apical ECM component as well.

### Morphogenesis of neurons and glia within an epithelium

Our results suggest a model in which epithelial properties of amphid neurons and glia play an integral role in their morphogenesis. First, amphid neurons and glia form an apical-basal polarized rosette shortly after the cells are born (Fan et al., 2018). Next, this rosette interacts with the developing epithelium of the skin and is carried to the nose (Fan et al., 2018). Finally, the cell bodies migrate away while their apical surfaces and cell junctions remain anchored at the nose (Heiman and Shaham, 2009). This anchoring requires apical ECM including DYF-7 and contributions from apically-localized FRM-2. In the absence of these factors, the developing tube-shaped glial pore ruptures at glial junctions and, although each of the individual cells retains its apical-basal polarity, the continuous lumen from sheath to socket is lost. These defects are comparable to phenotypes seen in the *C. elegans* excretory system or *Drosophila* tracheal tubes upon disruption of apical ECM (Jaźwińska et al., 2003; Wang et al., 2006; Stone et al., 2009; Mancuso et al., 2012; Gill et al., 2016; Forman-Rubinsky et al., 2017; Pu et al., 2017; Rosa et al., 2018). Thus, amphid dendrites are shaped by interactions within an epithelium composed of neurons, glia, and skin.

These features seem to be shared with other epithelial sense organs in *C. elegans* (CEP, IL, and OL). Intriguingly, retrograde extension has recently been reported in developing olfactory neurons in vertebrates. In zebrafish, olfactory placodes (which will give rise to olfactory epithelia) develop adjacent to the surface of the brain (Breau et al., 2017). Olfactory placode neurons attach nascent axons near the brain surface and then the cell bodies move away, extending axons behind them (Breau et al., 2017). The mechanism of anchoring in this case has not been described. However, given that much of our nervous system develops as a neuroepithelium – including sensory placodes, the cortex, and the cerebellum – it will be interesting to explore whether neuron-epithelial interactions might also help to shape dendrites and axons in other contexts.

## METHODS

### Strains and Plasmids

Strains were constructed in the N2 background and cultured under standard conditions (Brenner, 1974; Stiernagle, 2006). Strains, transgenes, and plasmids are listed in Supp. Tables S1-S3 respectively. All strains and plasmids are available upon request.

### Fluorescence microscopy and image processing

Animals were mounted on 2% agarose pads in water or M9 buffer (Sulston et al., 1983) with 0-50 mM sodium azide depending on developmental stage, and imaged using a Deltavision Core imaging system (Applied Precision) with UApo/340 40x 1.35NA, PlanApo 60x 1.42NA, and U-PlanApo 100x 1.4NA objectives (Olympus) and CoolSnap HQ2 camera. The amphid socket promoter (*grl-2*pro) yields bright expression in the excretory system that obscures the amphid features we wished to highlight in Fig. 4 and Supp. Fig. S1; we therefore used a fluorescence stereomicroscope to select rare mosaic animals that lacked excretory system expression to obtain the amphid images in these figures. Images were deconvolved using Softworx (Applied Precision) and maximum-brightness projections were obtained from contiguous optical sections using ImageJ. All projections used the same upper and lower limits across wavelengths, however, a thinner optical stack was often used in high-magnification images shown as insets. Due to large differences in signal intensity between neuronal and glial cell bodies and their thin processes, gamma settings were adjusted for the red signal in S1C in order to show the relevant structures clearly.

### Immunofluorescence

Embryos were fixed using a methanol-acetone procedure and stained with anti-AJM-1 primary antibody MH27 (Francis and Waterston, 1991) obtained from Developmental Studies Hybridoma Bank established at the University of Iowa by the NICHD/NIH) and sheep anti mouse DyLight649 secondary antibody (Jackson ImmunoResearch). Briefly, embryos were cracked by sandwiching them between a coverslip and Superfrost Plus slide (Fisher), then freezing them 5 min on a metal block on dry ice, and briskly removing the coverslip. The slide was placed in methanol at -20°C for 20 min followed by acetone at -20°C for 10 min, then washed in PBST (phosphate buffered saline with 0.1% Tween) and incubated in blocking solution (0.5% I-Block (ThermoFisher) in PBST) at room temperature for 1 h. Primary and secondary antibodies were used at 1:10 and 1:100 respectively in blocking solution. GFP fluorescence was preserved and visualized directly.

### Electron microscopy

Samples were subjected to high pressure freezing followed by freeze-substitution as described previously (Kolotuev et al., 2010). Samples were flat embedded, targeted and sectioned using the positional correlation and tight trimming approach (Kolotuev, 2014). This method facilitates identification of rare features in complex samples by using a precisely targeted region for sectioning. The sample was first viewed on a high-magnification optical microscope (Leica SPE confocal, 63x Leica oil immersion objective) and the distance between the first section and the expected location of the feature of interest was calculated. The sample was next mounted for ultramicrotome sectioning, and advancement of the sectioning was tracked following the feed count parameters of the ultramicrotome. 100nm sections were collected on the slot-formvar coated grids and observed with respect to the desired position using a JEOL JEM 1400 TEM microscope (JEOL, Japan).

### S2 cells

*Drosophila* Schneider (S2) cells (Invitrogen) were cultured and transfected with FuGene HD
(Roche) as described previously (Heiman and Shaham, 2009). Two days after transfection, cells and medium were harvested, applied in a small volume (~10 μl) directly to a coverslip without a slide, and imaged as described above.

### Genetic screen and mutant identification

Fourth larval stage (L4) animals of strain CHB99 (*oyIs44* V; *dyf-7(ns117)* X) were mutagenized using 70 mM ethyl methanesulfonate (EMS, Sigma) at 20°C for 4 h. Nonclonal F2 progeny were picked to 1000 individual plates. F3 populations were scored visually using a fluorescence stereomicroscope for increased (enhanced) or decreased (suppressed) penetrance of dendrite extension defects. Isolates corresponding to *hmn162* and *hmn169* exhibited 87% and 61% penetrance, respectively (n=100). Following a backcross to *dyf-7(ns117) kyIs136*[*str-2*pro:GFP] X males, cross-progeny exhibited 37% and 55% penetrance, respectively (n>150), suggesting a recessive mode of inheritance for *hmn162* and a dominant or partially dominant mode of inheritance for *hmn169*. In both cases, penetrance was similar between hermaphrodite and male cross-progeny, suggesting an absence of sex linkage. For mapping, 160 backcrossed F2 animals were picked to individual plates and the resulting F3 progeny were screened for enhanced penentrance of dendrite defects. 40 plates of animals with enhanced penetrance were identified, and these recombinants were pooled and subjected to genomic DNA extraction and whole-genome sequencing for one-step mapping (Doitsidou et al., 2010). The parental strains were also sequenced. Analysis with CloudMap (Minevich et al., 2012) identified a linked region on LG III including a nonsense mutation in *frm-2* (AAGTTTGTT[C>T]AGTGCAAGG).

### Dendrite length measurements

L4 animals were mounted and imaged as described above, and dendrite lengths were measured using the segmented line tool in ImageJ. The dendrite was traced from the point where it joins the cell body to the point where it ends at the nose, then normalized by the distance from the cell body to the nose to account for variance in the size of the animal.

## ACKNOWLEDGMENTS

We thank Shai Shaham, under whose support and guidance this project was initiated; Agnes
Burel and Marie Therese Lavault for assistance with electron microscopy; Elizabeth Lamkin and Karolina Mizeracka for plasmids; Jeff Simske and Verena Göbel for advice on AJM-1 and ERM-1 markers; Elisabeth Altendorfer and Monica Colaiacovo for assistance with immunofluorescence; Meera Sundaram, Zhirong Bao, and Li Fan for sharing unpublished data and generous advice; members of the Heiman lab for comments on the manuscript; and the *C. elegans* Genome Center and WormBase. This work was supported by an NSF Graduate Research Fellowship (IGM), March of Dimes Basil O’Connor Starter Scholar Award (MGH), and NIH R01GM108754 (MGH).

---

**Supplemental Table S1.** Strains used in this study

| **Strain** | **Genotype** | **Figure** |
| --- | --- | --- |
| CHB1566 | *hmnEx801* | 1D |
| CHB1612 | *hmnEx864* | 1E |
| CHB1665 | *hmnEx824* | 1F |
| CHB2721 | *hmnEx1535* | 1G |
| CHB2717 | *hmnEx1533* | 1H |
| CHB2719 | *hmnEx1534* | 1I |
| CHB440 | *dyf-7(ns119) X; hmnEx149* | 2B |
| CHB1939 | *hmnEx1105* | 2C |
| CHB436 | *oyIs44 V; dyf-7(ns119) X; hmnEx145* | 2D |
| CHB1530 | *hmnEx799* | 2E |
| CHB1557 | *hmnEx821* | 2F |
| CHB558 | *oyIs44 V; dyf-7(ns119) X; hmnEx216* | 2G |
| CHB1 | wild type | 3A |
| CHB157 | *dyf-7(ns119) X* | 3B |
| CHB1560 | *dyf-7(ns119) X; hmnEx824* | 4B,D |
| CHB2718 | *dyf-7(ns119) X; hmnEx1534* | 4C,E |
| CHB1556 | *dyf-7(ns119) X; hmnEx864* | 4F |
| CHB2720 | *dyf-7(ns119) X; hmnEx1535* | 4G |
| CHB2716 | *dyf-7(ns119) X; hmnEx1533* | 4H |
| CHB1743 | *oyIs44 V; hmnEx954* | 5A |
| CHB1741 | *oyIs44 V; hmnEx953* | 5A |
| CHB1745 | *oyIs44 V; hmnEx955* | 5A |
| CHB1739 | *oyIs44 V; hmnEx952* | 5A |
| CHB1742 | *oyIs44 V; dyf-7(ns119) X; hmnEx954* | 5B |
| CHB1740 | *oyIs44 V; dyf-7(ns119) X; hmnEx953* | 5B |
| CHB1744 | *oyIs44 V; dyf-7(ns119) X; hmnEx955* | 5B |
| CHB1738 | *oyIs44 V; dyf-7(ns119) X; hmnEx952* | 5B |
| CHB4 | *oyIs44 V* | 5C |
| CHB99 | *oyIs44 V; dyf-7(ns117) X* | 5C |
| CHB1211 | *frm-2(hmn162) III; oyIs44 V; dyf-7(ns117) X* | 5C |
| CHB2750 | *frm-2(hmn162) III; oyIs44 V; dyf-7(ns117) X; hmnEx1553* | 5C |
| CHB2950 | *hmnEx1667* | 5D |
| CHB2951 | *hmnEx1668* | 5E |
| CHB1477 | *hmnEx730* | S1A |
| CHB1343 | *hmnEx660* | S1B |
| CHB1475 | *hmnEx756* | S1C |
| CHB370 | *oyIs44 V; dyf-7(ns119) X; hmnEx97* | S2A |
| CHB443 | *oyIs44 V; dyf-7(ns119) X; hmnEx151* | S2A |
| CHB436 | *oyIs44 V; dyf-7(ns119) X; hmnEx145* | S2A |
| CHB1613 | *hmnEx865* | S2B |
| CHB1615 | *hmnEx867* | S2C |

**Supplemental Table S2.**
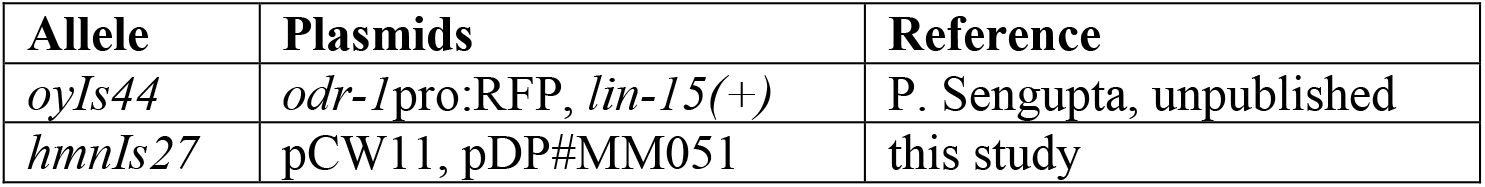
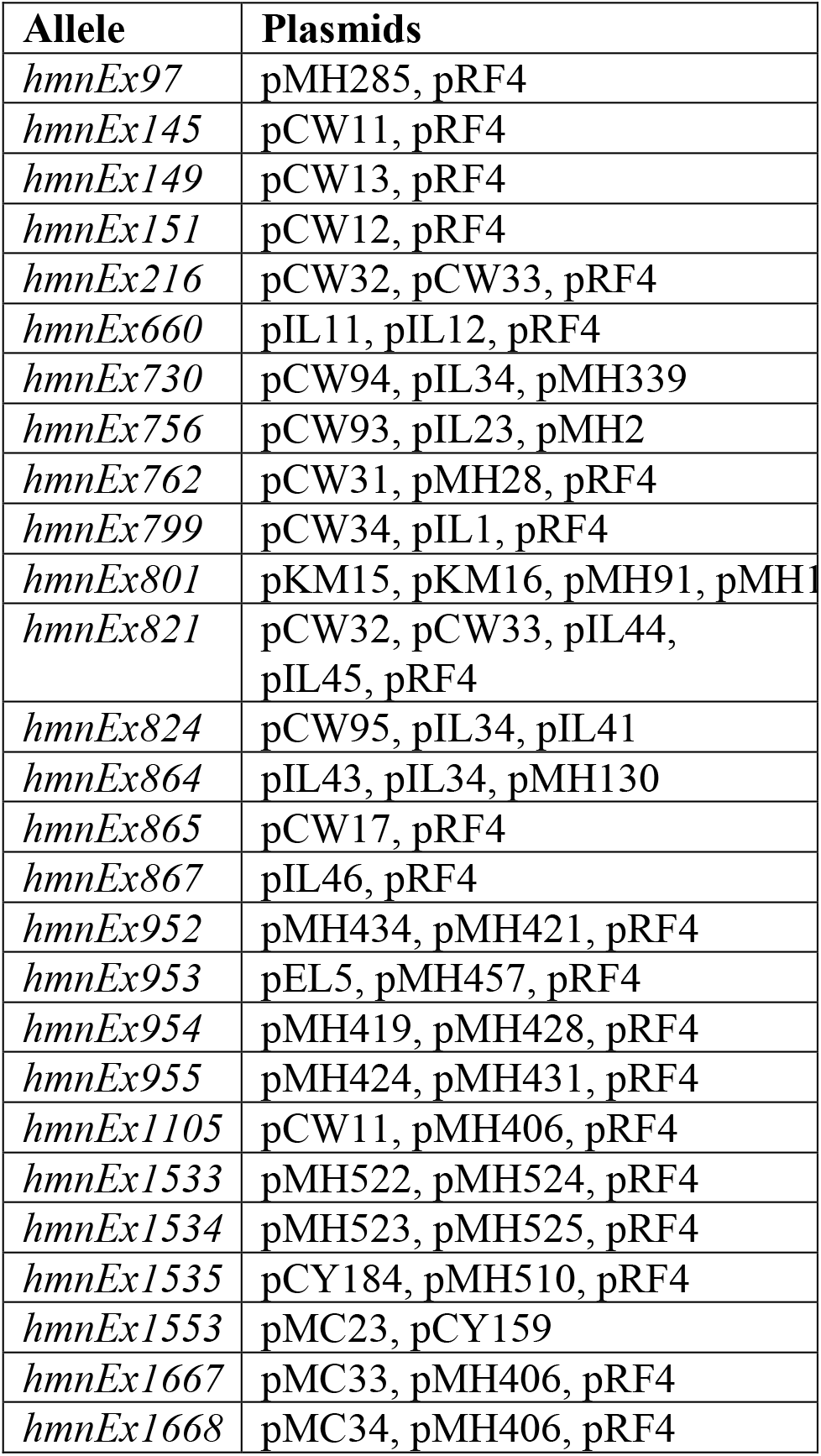
Transgenes used in this study

**Supplemental Table S3.**
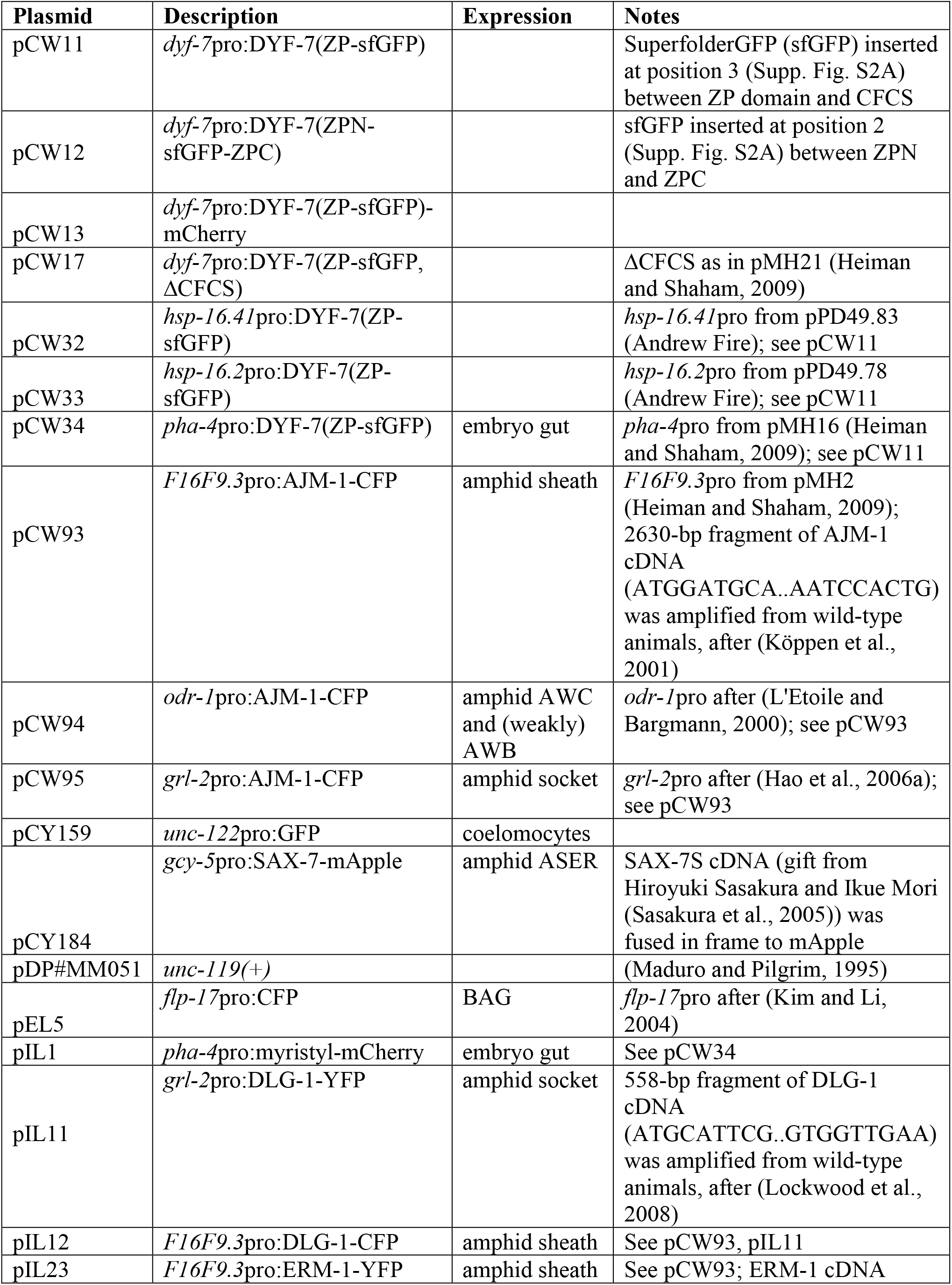
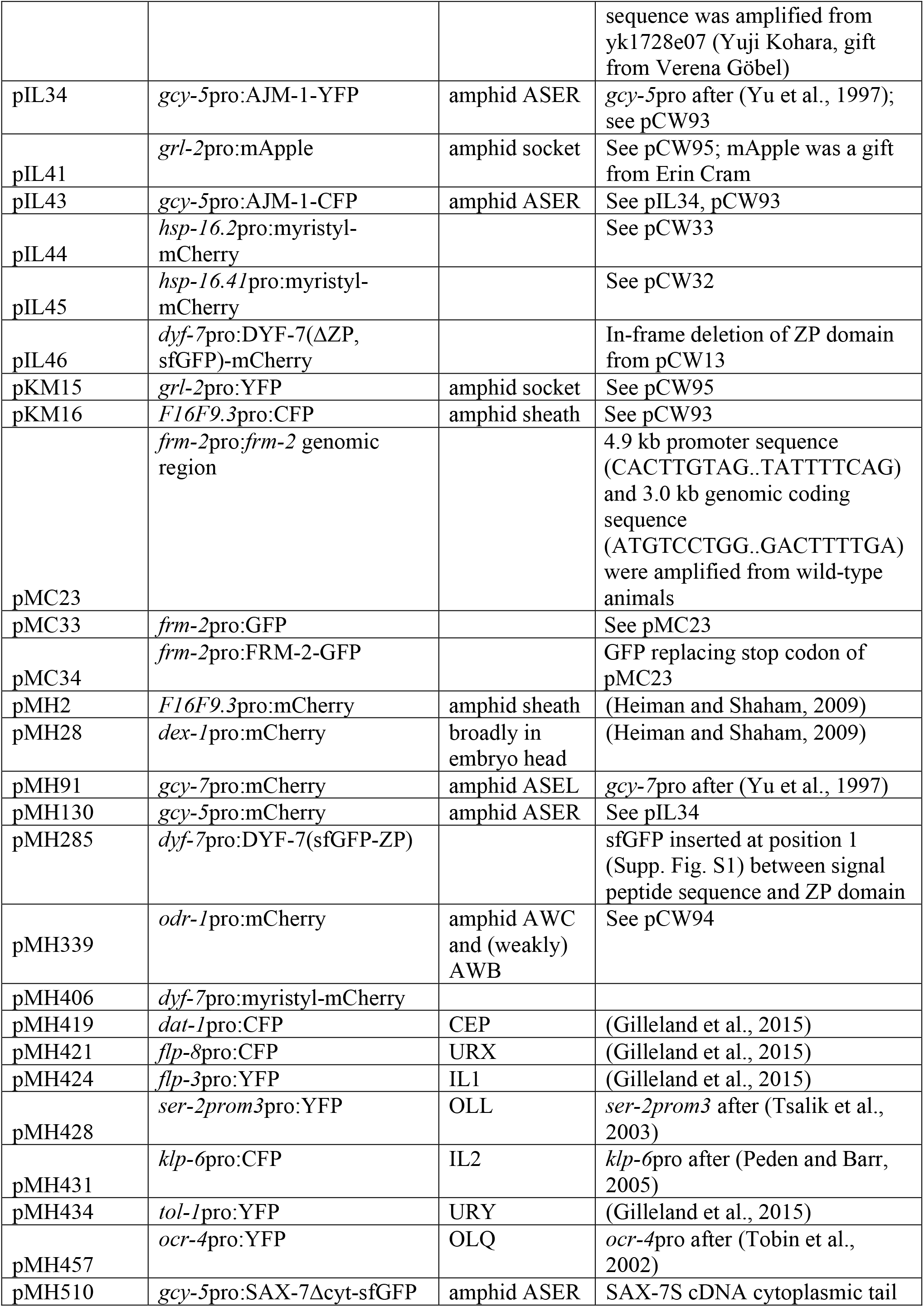
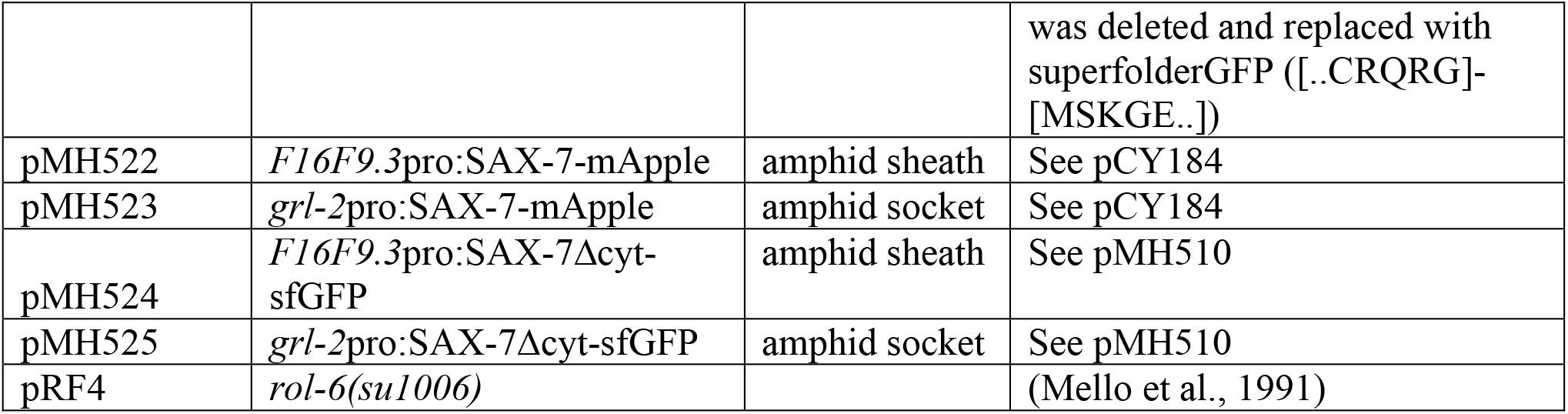
Plasmids used in this study

**Supporting Figure S1.**
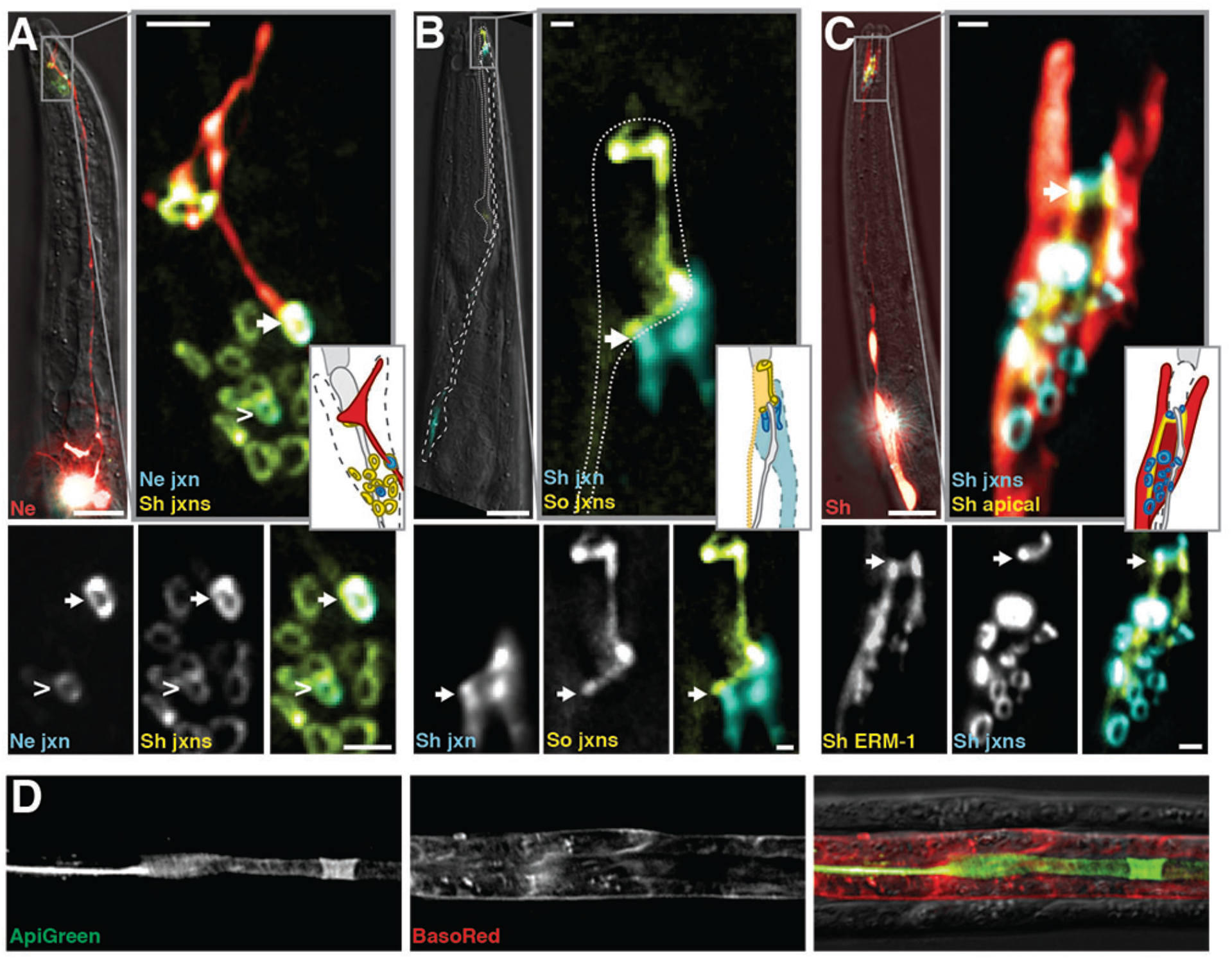
Markers for tight junctions and apical surfaces. (A, B) Overlap of neuron:sheath junctions as in Fig. 1, except (A) showing amphid neurons AWC (arrow) and AWB (carat) instead of ASE (Ne, AWC neuron (red, *odr-1*pro:RFP, only AWC is shown as AWB expression is dimmer and not visible here); Ne jxn, neuron junctions (blue, *odr-1*pro:AJM-1-CFP); Sh jxns, sheath junctions (yellow, *F16F9.3*pro:AJM-1-YFP)), and (B) showing the tight junction marker DLG-1 instead of AJM-1 (Sh jxn, sheath junctions (blue, *F16F9.3*pro:AJM-1-CFP, only sheath:socket junction is shown as sheath:neuron junctions are outside the region of interest); So jxn, socket glial junctions (yellow, *grl-2*pro:DLG-1-YFP)). (C) The apical cytoskelal-associated protein ERM-1 localizes to the outward-facing surface of the sheath (arrow). Sh, sheath glial cell (red, *F16F9.3*pro:mCherry); Sh jxns, sheath junctions (blue, *F16F9.3*pro:AJM-1-CFP); Sh ERM-1 (yellow, *F16F9.3*pro:ERM-1-YFP). (D) Expression of apical marker ApiGreen and basolateral marker BasoRed in gut, a well-studied model epithelium. The lumen (apical) surface of the gut exhibits localization of ApiGreen and exclusion of BasoRed. Scale bars: 10μm, main panels; 1μm, magnified insets.

**Supporting Figure S2.**
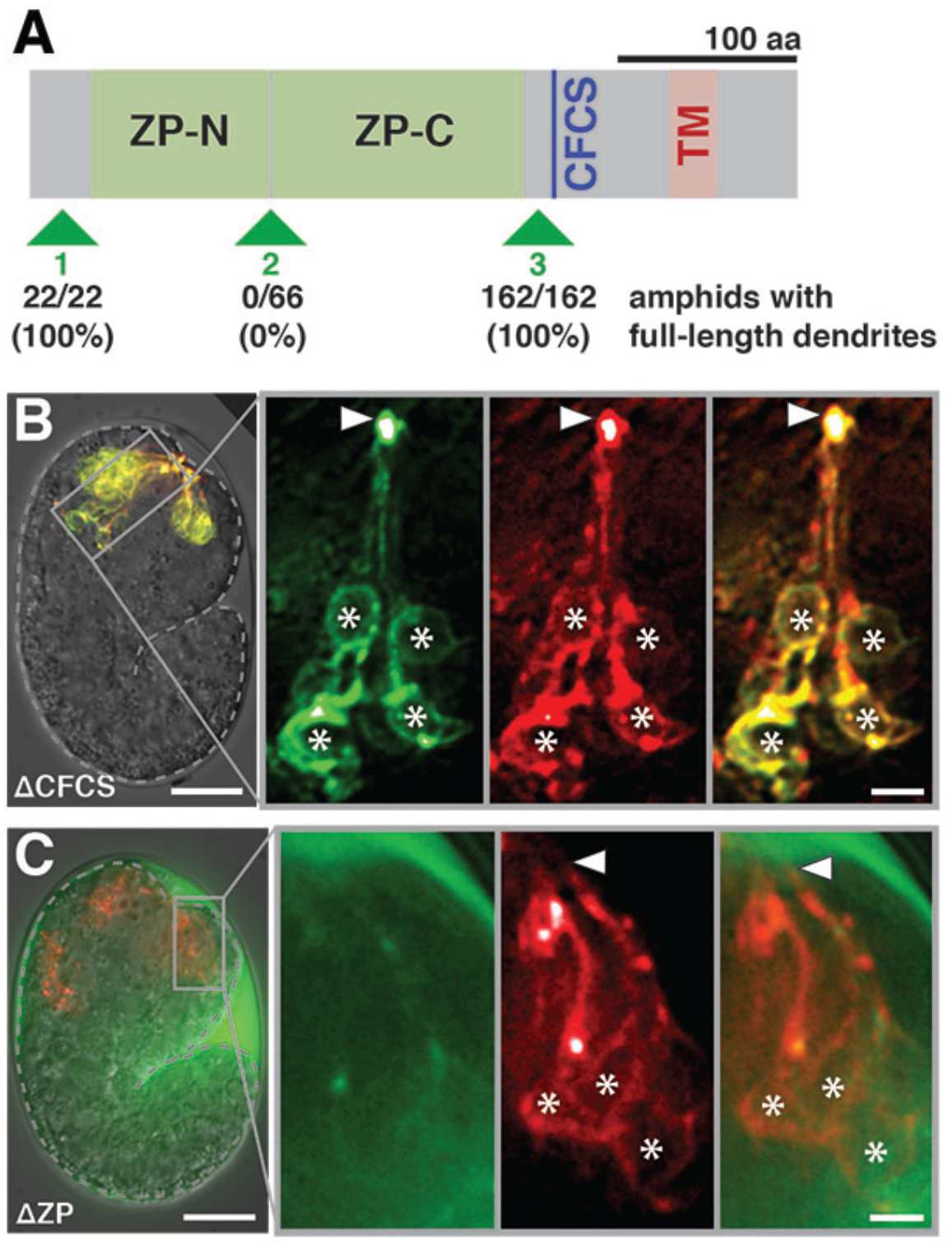
Localization of DYF-7 *in vivo*. (A) SuperfolderGFP coding sequence was inserted into the DYF-7 cDNA at positions 1, 2, or 3 as indicated (pMH285, pCW12, and pCW11, respectively; Table S1). These constructs were introduced into a strain bearing *dyf-7(ns119)* and an amphid neuron marker (AWC, *odr*-*1*pro:RFP) and scored for rescue of dendrite extension defects. CFCS, consensus furin cleavage site; TM, transmembrane segment. (B,C) Live, intact embryos expressing sfGFP-DYF-7-mCherry in sensory neurons (*dyf-7*pro) at the time of dendrite extension, as in Fig. 2, but with constructs that are (B) lacking the CFCS (ΔCFCS) or (C) ZP domain (ΔZP). sfGFP, green; mCherry, red; arrowheads, dendrite tips; asterisks, neuron cell bodies. Scale bars: 10μm, main panels; 2μm, magnified insets.

